# Real-Time Multi-Position and Multi-ROI Tracking with LiLiTTool for Smart Light-Sheet Microscopy in Growing Samples

**DOI:** 10.64898/2026.04.03.715281

**Authors:** Clément Helsens, Frédéric Pili, Emanuel Vasquez, Florian Aymanns, Jean-Yves Tinevez, Edward Andò, Andrew Oates

**Affiliations:** Timing, Oscillators, Patterns Lab, Institute of Bioengineering, Swiss Federal Institute of Technology Lausanne, EPFL; Center for Imaging, Swiss Federal Institute of Technology Lausanne, EPFL; Institut Pasteur, Université Paris Cité, Image Analysis Hub, F-75015 Paris, France

**Keywords:** smart microscopy, light-sheet microscopy, real-time tracking, offline tracking, cotracker3

## Abstract

Long-term live imaging of growing samples with light-sheet fluorescence microscopy provides unique insights into development, but morphogenesis often displaces features of interest outside the microscope’s field of view (FOV), calling for automated methods to track these features and update the microscope’s FOV in real time. Existing solutions, which typically rely on local or global intensity distributions, struggle to follow specific features robustly throughout morphogenesis, leading to truncated or incomplete datasets. Here, we present a light-sheet live tracking tool (LiLiTTool) that maintains user-defined regions of interest (ROI) within the FOV throughout extended imaging sessions. LiLiTTool uses Cotracker3, a state-of-the art deep learning–based motion predictor, augmented by sensor fusion with a trained object-detector. This enables robust compensation for drift, rotation, and deformation during morphogenesis, while meeting the timing constraints of live acquisition. We validated LiLiTool by integrating with the Viventis LS1 microscope, achieving sub-second processing and stable tracking of growing zebrafish embryos over many hours. LiLiTTool supports multi-ROI tracking in 3D, enabling simultaneous monitoring of multiple features within the same embryo and in multiple embryos during a single acquisition. LiLiTTool is modular and openly available on GitHub and as a napari plugin for post-acquisition tracking. By enabling precise, adaptive, and scalable real-time imaging, LiLiTTool advances smart microscopy approaches and provides the developmental biology community with a practical tool for capturing reliable spatio-temporal information in growing embryos or other morphogenetic systems.

## Introduction

Time-lapse microscopy is a key method in the study of dynamics in developmental biology, as well as in other multicellular morphogenetic systems such as organoids. Light-sheet fluorescence microscopy (LSFM) has emerged in the last two decades as a method of choice for fast, low-toxicity volumetric imaging, allowing morphogenetic processes to be studied at cellular and sub-cellular resolution over hours or days (1, 2). However, over these time-scales samples may rotate, deform, elongate and grow, leading to substantial changes in their position and geometry within the imaging volume. Consequently, the anatomical feature of interest and its constituent cells define a region of interest (ROI) that may move out of the field of view (FOV) during imaging, rendering the resulting data unsuitable for downstream analysis. Given the substantial time, effort, and cost required to prepare and image biological samples, this limitation remains a major challenge for experimental studies (3, 4).

As an alternative to a spatially defined feature of interest, the entire specimen can be imaged throughout development using a microscope with a sufficiently large FOV, or a systematic tiling approach. Such *in toto* imaging may be a goal in its own right (2, 5, 6); features of interest can then be extracted from the recording for closer study offline. However, current microscopes with these capacities are complex, expensive and custom-built by expert teams and their recording sizes are in the tens of terabyte range. Moreover, the problems of drift and motion are in practice still present, either in the need to adapt acquisition parameters during recording, or to achieve ROI tracking after the completion of the experiment (7, 8). A recent review (9) discusses the challenges of large, multidimensional light-sheet imaging datasets, the inadequacy of manual intervention, and the need for continued development of automated imaging pipelines that update microscope parameters in response to changes in the image of the sample, an approach termed smart microscopy (10–15).

A specific application of smart microscopy aims to automatically reposition the microscope stage in X, Y, and Z during closed-loop acquisition, so as to follow a region of interest (ROI) and maintain it at a fixed location within the field of view (FOV) as the sample develops and deforms. This requires integrating a feedback loop into the acquisition procedure, in which image analysis is used to estimate ROI motion within the FOV and drive compensatory stage movements. We refer to this process as ROI tracking throughout this article.

For many laboratories, ROI tracking remains a key requirement to enable long-term imaging–based discovery. The developing zebrafish embryo illustrates this challenge: although it is well suited to time-lapse imaging due to its optical transparency, its rapid morphogenesis leads to pronounced tissue deformation and multi-fold elongation of the body axis. In our work on somitogenesis, these dynamics consistently caused the anatomical feature of interest - the tailbud and presomitic mesoderm - to drift out of the FOV, resulting in failed acquisitions.

To address this, we initially relied on an intensity-based center-of-mass tracking strategy applied to a localized, tissue-restricted fluorescent signal, allowing the motion of the zebrafish tail to be coupled automatically to the microscope stage position (16, 17). However, this approach proved vulnerable to several sources of error. Global or local intensity fluctuations biased the center-of-mass estimate; transient bright artifacts within the FOV could attract the estimator away from the target structure; and developmental changes in tissue shape altered the spatial distribution of the signal. Projection effects and background fluorescence further reduced robustness.

Beyond these technical limitations, intensity-based approaches are intrinsically constrained by the availability of suitable transgenic reporters or probes that selectively label the structure of interest, limiting their general applicability. More generally, methods based on center-of-mass estimation or local correlation are prone to failure in the presence of local deformation or non-uniform motion (6–8). Despite these efforts, more than one third of our long-term acquisitions were ultimately unsuccessful.

In this work, we present LiLiTTool, a ROI tracking framework that combines modern deep learning and classical image registration with sensor fusion methods, tightly integrated with microscope hardware. By coupling anatomically specific tracking—focusing on user-defined morphological features—with real-time adjustment of the microscope stage, our approach ensures stable long-term imaging of features of interest within a restricted FOV in moving embryos without specific labels. Although LiLiTTool was optimized using growing and elongating zebrafish embryos as a test system, we further validated its performance on additional types of biological specimens, confirming its general applicability across different organisms, spatial scales, and imaging modalities. LiLiTTool was successfully integrated into the Viventis LS1 LSFM environment. However, the software was designed to be hardware-agnostic and can, in principle, be coupled to any microscope system, provided that the acquisition software exposes the required trigger signals and stage control commands through a suitable application programming interface (API).

## Results

### LiLiTTool overview - design choices and implementation

To address the challenges of imaging developing tissues, we require an efficient image analysis algorithm capable of recognizing and tracking a ROI as it moves, grows, and deforms within the microscope FOV. ROI tracking must operate directly on the imaging channel of interest, regardless of the features it contains, in order to avoid reliance on a dedicated fluorescent marker.

Traditional image analysis approaches typically rely either on the explicit detection of predefined features or on pixel-wise optimization of standard similarity metrics, such as cross-correlation. Although these methods are capable of impressive feats for tracking, the complexity of motion in morphogenetic systems, together with the variability of tissue features commonly imaged in the life sciences, limits their robustness and general applicability for our purposes.

Deep learning (DL) approaches, by contrast, have demonstrated a strong capacity to leverage diverse image content. Modern implementations exploit dedicated hardware, such as graphical processing units (GPUs), enabling high computational performance compatible with robust real-time tracking. Among DL algorithms, a class of neural networks performs data-driven feature extraction by learning complex spatial and temporal representations directly from data. These models excel in challenging tracking tasks that require recognition of higher-level patterns and preservation of temporal consistency over extended time periods.

In particular, the CoTracker3 model (18, 19) has recently emerged as a high-performance point-tracking algorithm with broad applicability. CoTracker3 is a transformer-based neural network designed to track arbitrary points in 2D video sequences, combining convolutional feature extraction with transformer-based temporal modeling to robustly follow multiple points over time. In this work, we adopted CoTracker3 as the computational core of LiLiTTool. LiLiTTool’s core codebase is architecturally centered on this point-tracking algorithm and is complemented by modules specifically designed for biological imaging. These additional components enhance robustness against the complex morphological changes and dynamic evolution characteristic of developing samples.

Implementation details of LiLiTTool are provided in the Material and Methods section. The overall acquisition procedure including the configuration of LiLiTTool and its internal software-hardware interactions are show on Fig. 1.

**Fig. 1.**
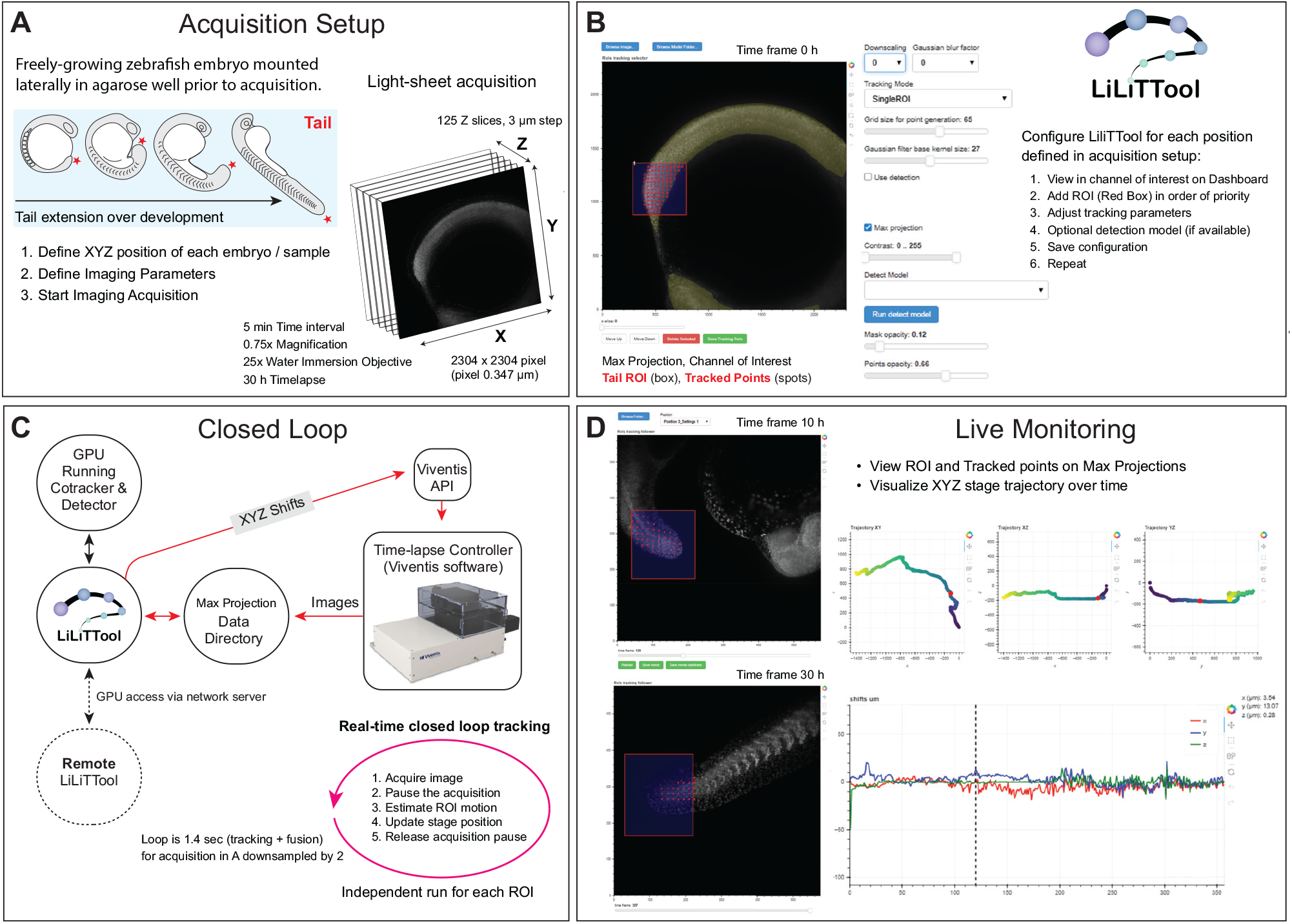
Overview of the experimental setup, LiLiTTool configuration, and real-time monitoring during smart microscopy acquisition. **(A) Experimental preparation and imaging conditions**. A freely-growing zebrafish embryo is mounted in an agarose well in E3 medium prior to acquisition. Imaging is performed on a Viventis LS1 light-sheet microscope using typical time-lapse parameters for long-term developmental imaging. These include volumetric stacks acquired at regular time intervals, with multiple imaging channels, defined stack dimensions, and high-speed image saving to disk. Image data are stored in a high-throughput file format and written continuously to a monitored directory. During initialization, the structure of interest to be tracked is positioned near the center of the microscope field of view (FOV) to allow symmetric stage corrections in X, Y, and Z during acquisition. **(B) LiLiTTool configuration interface.** Prior to acquisition, tracking parameters are configured through an interactive dashboard. For each imaging position, the user defines the region(s) of interest (ROIs) to be tracked and selects the desired tracking mode. Additional parameters include the density of query points used by the tracking algorithm, the level of Gaussian blurring applied to the image prior to query point generation, whether blurring is applied during tracking to mask internal texture movement, and whether object detection should be used for sensor fusion. These parameters can be adjusted independently for each position and are saved as part of the experiment configuration before starting the time-lapse. **(C) Software–hardware interaction and live monitoring during acquisition**. The diagram illustrates the communication between the Viventis LS1 light-sheet microscope, the Viventis time-lapse controller, LiLiTTool, and local or remote GPU resources. After each time point is acquired at a given imaging position, the microscope emits a trigger signal that initiates LiLiTTool processing. After the microscope acquisition loop is paused, LiLiTTool then performs tracking, either locally or on a remote GPU, computes corrected stage coordinates (X, Y, Z), and transmits the corresponding stage-shift commands back to the microscope. Once the correction is applied, and the paused released, acquisition proceeds to the next position. This process is repeated for all positions at each time point throughout the time-lapse experiment, while a monitoring dashboard allows users to visualize the tracking status in real time. **(D) Real-time monitoring dashboard**. A dedicated monitoring interface allows users to follow the tracking process interactively during acquisition. The dashboard displays the incoming images, the current ROI positions, and the tracked query points, enabling real-time assessment of tracking performance and system status throughout the experiment.

#### LiLiTTool case study

The case study focused on the tip of the zebrafish embryo’s tail (tailbud), a dynamically evolving anatomical structure that undergoes substantial displacement during embryonic development. We performed long-term light-sheet imaging of zebrafish embryos mounted laterally with the yolk nestled in an agarose well. All experiments were conducted using an XY light-sheet acquisition scheme with multiple Z slices acquired at regular time intervals (typically 5 minutes). Over a typical 15 to 30-hour acquisition, the cumulative displacement of the tail tip along the lateral plane (XY) exceeded twice the lateral field of view of the microscope (approximately 700 *µ*m). This highlights the necessity for real-time, structure-specific tracking to maintain the feature of interest within the field of view (Fig. 2 top row and supplemental movie 1).

**Fig. 2.**
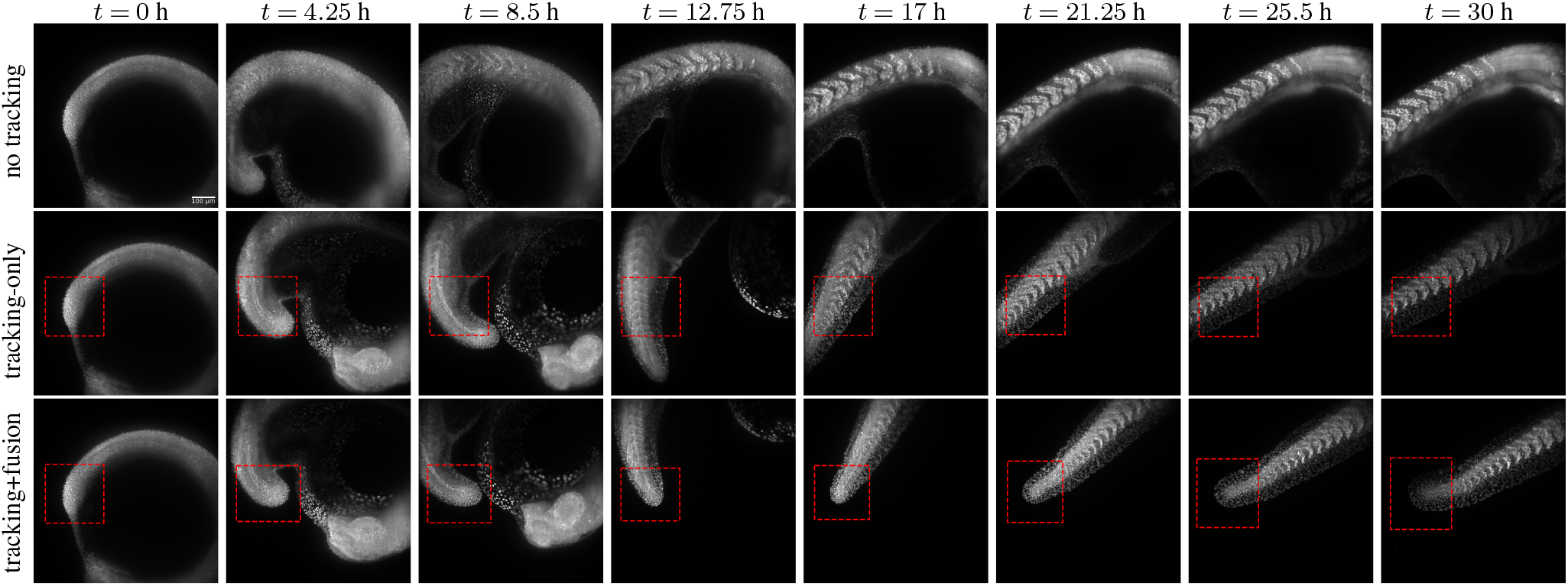
Effect of LiLiTTool real-time tracking on maintaining the region of interest during long-term imaging. Comparison of a 30-hour light-sheet time-lapse acquisition of the same zebrafish embryo performed without tracking (top row), with *tracking-only* (middle row), and with *tracking+fusion* (bottom row) of the tail tip using LiLiTTool. Imaging carried out under identical conditions on the LS1 Live microscope. **Without tracking**, natural embryo growth and motion cause the tail to progressively drift out of the field of view, leading to partial or complete loss of the target region shortly after the start of the acquisition. **With *tracking-only*** enabled, incoming images are continuously analyzed by LiLiTTool, and the estimated tail motion is compensated by active repositioning of the microscope stage. This maintains the tail region within the field of view for an extended period. However, the target is eventually lost because the tracker follows internal cellular structures rather than the global anatomical structure. **With detection enabled (*tracking+fusion*)**, LiLiTTool identifies the tail tip once it detaches from the yolk and subsequently tracks it accurately for the entire duration of the experiment. The red box in the middle and bottom rows indicates the region of interest (ROI) selected for tracking by LiLiTTool.

Below, we describe the two primary strategies in LiLiTTool to overcome the loss of the tail tip from the FOV: *tracking-only* and *tracking+fusion*. In our case, with *tracking-only*, complex internal movements can result in the tail tip being eventually lost from the FOV (Fig. 2 middle row and supplemental movie 2), but with *tracking+fusion* the tail tip is retained for the entire duration of the movie (Fig. 2 bottom row and supplemental movie 3). Using this configuration, we successfully tracked and imaged multiple embryos simultaneously for up to 30 hours, maintaining stable positioning of the target regions on the FOV throughout the acquisition.

#### LiLiTTool core algorithmic implementation

After acquisition of the first frame Fig. 1a, the user defines a ROI on the image Fig. 1b. XY tracking is then performed on a maximum-intensity projection of the acquired volume Fig. 1c.

To generate query points for tracking, the projected image is first smoothed with a Gaussian filter to reduce local intensity fluctuations and stabilize the signal prior to segmentation. Otsu’s thresholding method (20) is then applied to produce a binary mask of the tissue region. A uniform grid of candidate points is generated across the entire image, and points located outside the segmented tissue mask are discarded. The final set of query points corresponds to the intersection between the binary mask and the user-defined ROI, ensuring that tracking is confined to biologically relevant regions (red points in Fig 1 and supplementary material Figure 1).

The selected query points are provided as input to Co-Tracker3, which processes subsequent projected frames to estimate their updated positions over time (see supplementary material Figure 1).

The displacement of these points is used to compute an updated ROI position (see Material and Methods), which in turn drives compensatory stage movements in X and Y so that the ROI remains at its initial location within the FOV in the next frame. Sample movements in X-Y are predicted using CoTracker3 with a default sliding window of ten frames.

Displacement along the Z axis is estimated independently using a classical image registration approach based on an intensity-weighted center of mass computed within the projected ROI (see Materials and Methods). This procedure (X-Y then Z) is iterated throughout the acquisition.

This design reflects typical sample mounting strategies for volumetric imaging, where live specimens are oriented such that morphological complexity and displacement are more pronounced in the XY plane rather than along Z. For example, when imaging the zebrafish embryo tail, embryos are mounted laterally so that the tail extends primarily within the XY imaging plane. Motion along Z is generally of lower amplitude and involves simpler structural geometry, making it suitable for estimation with a shape-dependent registration method. This strategy—tracking the ROI in XY using CoTracker3 and estimating Z displacement via image registration—is referred to as *tracking-only* in LiLiTTool throughout the remainder of this article.

In multi-position acquisitions, the degree of Gaussian smoothing and the subsequent density of query points can be configured independently for each position. LiLiTTool also provides several tracking modes to support the simultaneous tracking of multiple ROIs at a given position (see Material and Methods).

#### Sensor fusion and texture filtering

Despite its power, the *tracking-only* strategy can present two limitations when applied to developing samples, which can be addressed by additional steps presented below.

First, because the ROI position at each time point is updated exclusively through motion prediction, small estimation errors may accumulate over time, resulting in gradual drift. This phenomenon is intensified when competing motions occur within the sample. In the zebrafish embryo, for example, the tail tip and the differentiating progenitor cells that emerge from the tailbud move in partially opposing directions. As a point-based tracker, CoTracker3 is naturally attracted to localized features, such as those generated by individual cells. Consequently, it may preferentially follow local cellular motion rather than the global displacement of the anatomical structure of interest (Fig. 2 middle and supplemental movie 2).

Second, the ROI defined at initialization remains fixed in size and shape throughout the time-lapse. As the embryo grows and deforms, initial query-points within the ROI drift apart. As the updated ROI fits these new points, it can become misaligned with the underlying anatomical feature of interest, leading to suboptimal placement of new query points and reduced tracking accuracy.

To address the first limitation, we implemented an optional Gaussian blurring step to reduce the influence of high-frequency texture motion on CoTracker3. This preprocessing approach performs well when the feature of interest is already clearly identifiable at the start of acquisition, *e*.*g*. when the tail has already detached from the yolk. Its effectiveness decreases, however, when the feature of interest is not yet distinctly present within the initial ROI, as in earlier developmental stages, *e*.*g*. when the tailbud remains flattened against the yolk. This approach of using texture filtering prior to tracking is termed *blurred-tracking*.

Both limitations—internal textural motion and fixed ROI size—can be mitigated using a sensor fusion strategy that augments motion-based tracking estimates with information derived from an independent algorithmic source. We therefore introduced object detection as a complementary source of positional information. A dedicated detection model, trained to recognize a specific morphological feature, is integrated into LiLiTTool. This detection provides external observations that are combined with the tracking-only estimates through a sensor fusion framework (21–24). These externally derived updates enhance the accuracy and stability of ROI tracking.

For zebrafish tail development experiments, we trained a model to detect the tail tip. At early developmental stages, the tail tip remains attached to the yolk, and CoTracker3 provides the primary source of positional updates. Once the tail tip detaches and becomes morphologically distinct, it is recognized by the object detector, and its positional estimates are fused with those of CoTracker3. This integration corrects drift arising from changes in image texture and allows dynamic adjustment of the ROI to the evolving anatomical feature. We implemented the detection component using Faster R-CNN (25, 26) trained on ground-truth annotated datasets of the zebrafish tail tip. The resulting combined strategy retains the tip of the tail in the FOV and is referred to as *tracking+fusion* in LiLiTTool and is shown in Fig. 2 bottom row (and supplemental movie 3).

In summary, when internal textured motion is minimal, CoTracker3 alone is typically sufficient for accurate tracking, irrespective of image blurring. When textured motion is present but can be attenuated within the ROI, Gaussian blurring may be adequate to maintain reliable tracking of larger features. However, when the initial configuration or developmental dynamics prevent these simpler solutions, sensor fusion with an appropriate detection model provides superior robustness and accuracy.

### Comparison of LiLiTTool to human expert

To quantify the tracking performance of LiLiTTool, we performed an *in silico* experiment by re-tracking an existing time-lapse movie offline. A movie in which the tail tip had already detached from the yolk and was reliably tracked during acquisition was selected. A human expert was asked to manually define the ROI corresponding to the tail tip every five frames throughout the sequence. These annotations were used as a reference trajectory for comparison.

We then reprocessed the same movie using three different configurations of LiLiTTool: *tracking-only* (green), *blurred-tracking* (yellow), and *tracking+fusion* without blurring (blue). The positions of the ROIs estimated by each configuration were compared to the expert-defined reference (red). As shown in Fig. 3, both the blurred *tracking-only* configuration and the *tracking+fusion* approach closely follow the trajectory defined by the expert tracker, whereas the *tracking-only* configuration without blurring progressively deviates from the target structure due to the influence of internal cellular motion.

**Fig. 3.**
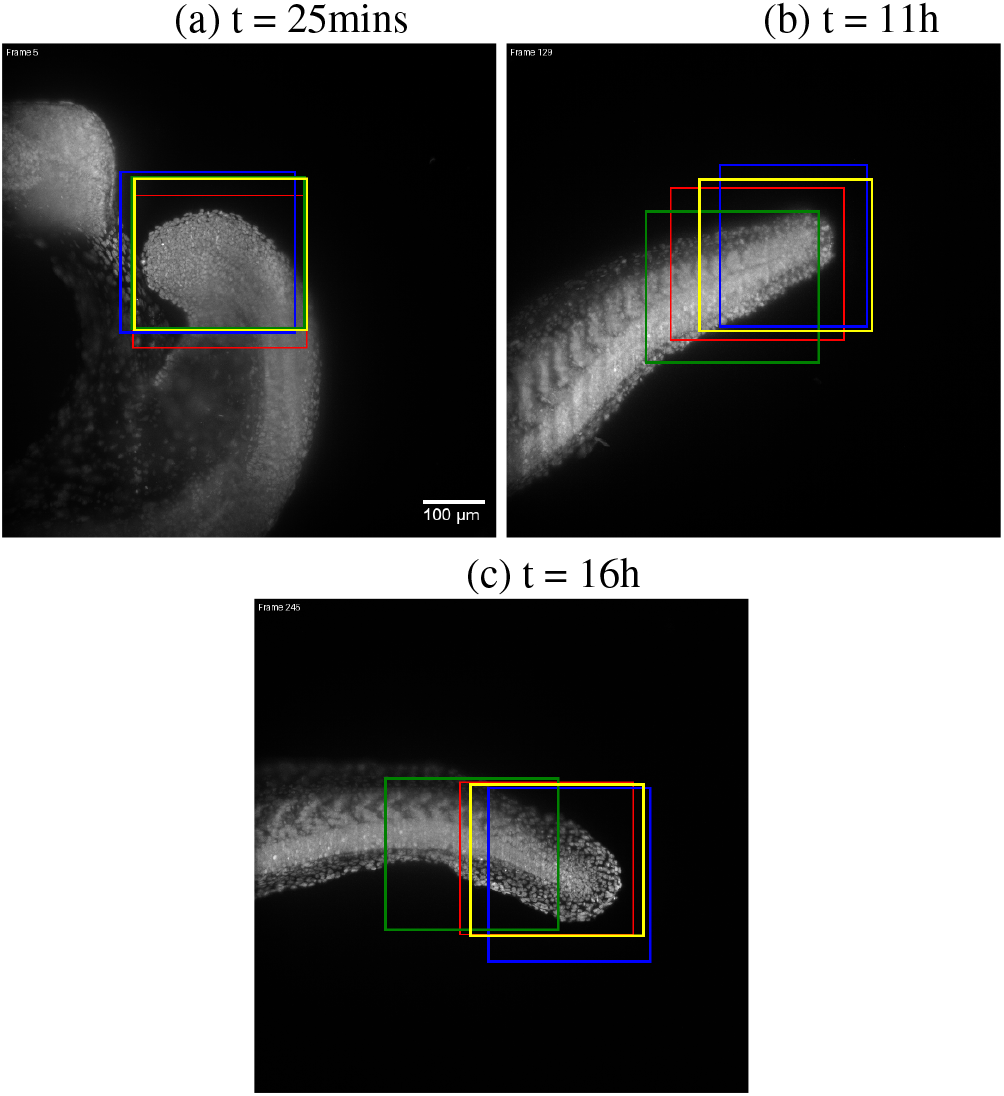
Evaluation of LiLiTTool tracking performance in an *in silico* re-tracking experiment. Comparison of different tracking strategies processed offline using a previously acquired time-lapse movie of a zebrafish embryo tail tip. An ROI containing the tail tip defined every five frames by a human expert provides a reference trajectory (red boxes). The movie is re-tracked using three LiLiTTool configurations: *tracking-only* (green), *blurred-tracking* (yellow), and *tracking+fusion* without blurring (blue). Three representative time points are shown: early (t = 25 min, (a)), mid (t = 11 h, (b)), and late (t = 16 h, (c)). Both *blurred-tracking* and *tracking+fusion* approaches closely follow the expert-defined trajectory, whereas *tracking-only* progressively deviates from the target structure.

### Successful tracking examples in more difficult conditions

We imaged zebrafish embryos in low fluorescence conditions, relying only on the auto-fluorescence of the fish. Despite the low signal intensity and reduced signal-to-background contrast, the tracking tool was still able to accurately follow the selected region of interest (see Movie 5 in supplementary material).

We also tested LiLiTTool during a heat-shock experiment in which the embryo undergoes a rapid temperature increase. This lead to contractions of the body axis and sudden large displacements of the tail. Despite these abrupt movements and tail deformation, we were able to track the selected ROI (see Movie 6 in supplementary material).

We further assessed the robustness of LiLiTTool in situations where multiple biological structures are present in the imaging field. In this experiment, two zebrafish embryos were simultaneously present within the same FOV, while LiLiT-Tool was configured to track the tail tip of only one embryo. Despite the additional structures and motion introduced by the second embryo, LiLiTTool maintained stable tracking of the selected target throughout the acquisition, demonstrating its ability to remain focused on the user-defined ROI even in complex imaging environments (see Movie 7 in supplementary material).

### Generalization to other organisms and 2D datasets

To further evaluate the generalizability of the proposed tracking framework, we conducted experiments on a range of 2D datasets featuring diverse biological specimens and imaging modalities using *tracking-only*. These included (i) Tardigrades (fluorescent, see supplementary material Movie 8), (ii) *Caenorhabditis elegans* embryos imaged during mitosis from unpublished datasets acquired in other laboratories at EPFL (bright field, see supplementary material Movie 9), and (iii) human hepatocarcinoma-derived (HepG2) cells from the Cell Tracking Challenge benchmark datasets (27) (fluorescent, see supplementary material Movie 10).

In the Tardigrade dataset, the framework was used to track the position of the animal’s head across time, maintaining stability despite strong deformations and non-rigid movements of the organism. In the *C. elegans* experiments, the method accurately followed the fluorescently-labelled spindle poles during cell division, capturing their fast and non-linear trajectories within a crowded cellular environment. Finally, on the Cell Tracking Challenge dataset, the system successfully tracked 17 selected individual HepG2 cells independently throughout the sequence, consistently preserving cell identities and trajectories over time.

Across all datasets, LiLiTTool *tracking-only* mode demonstrated strong robustness and adaptability without the need for organism-specific retraining. These results showed the method’s capacity to generalize across systems and scales, imaging setups, and motion dynamics, confirming its potential applicability to a broad range of biological imaging contexts.

### LiLiTTool software implementation

To facilitate the use of our method, it is shared as a stand-alone software package that enables both real-time and offline tracking (https://github.com/EPFL-TOP/lightsheet-live-tracking-tool). The package is modular and easily adaptable for use with other microscopy systems. In addition to real-time tracking, it supports offline processing of ROIs, forward and backward point tracking, and provides an integrated Napari (28) plugin for visualization and user interaction (https://github.com/EPFL-TOP/cotracker-napari-plugin).

When local GPU resources are insufficient or should remain dedicated to the ongoing acquisition, inference tasks can be seamlessly offloaded to remote GPUs using the `imaging-server-kit` framework (https://github.com/Imaging-Server-Kit/imaging-server-kit). Built-in fail-safes ensure that acquisitions continue uninterrupted, in the event of inference failure.

The LiLiTTool software was directly integrated with the Viventis LS1 Live microscope, demonstrating sub-second processing times and stable long-term tracking up to 30 hours. In addition to single structures, the software also supports multi-ROI tracking, enabling simultaneous monitoring of multiple anatomical regions within the same embryo or tracking of multiple embryo positions during the same acquisition.

### LiLiTTool tracking accuracy assessment

Tracking accuracy was evaluated using a self-consistency–based strategy in the absence of a unique ground truth trajectory. Because maintaining a specific anatomical feature within the field of view does not admit a single correct solution, we assessed performance by re-tracking previously acquired datasets after introducing known artificial displacements. The tracking pipeline reliably recovered these imposed shifts, yielding residual errors centered close to zero with low variability along both lateral axes (mean residuals of approximately ± 0.2 pixels and standard deviations below 2.1 pixels). These residuals are small relative to both the microscope field of view and the characteristic scale of tail motion, indicating minimal systematic bias and stable performance over long acquisitions (Fig. 4). Together, these results demonstrate that the proposed tracking approach accurately compensates for lateral motion and remains robust under controlled perturbations, even in the absence of explicit ground truth annotations. For a more detailed explanation of the method, see the dedicated sub-section in the materials and methods section.

**Fig. 4.**
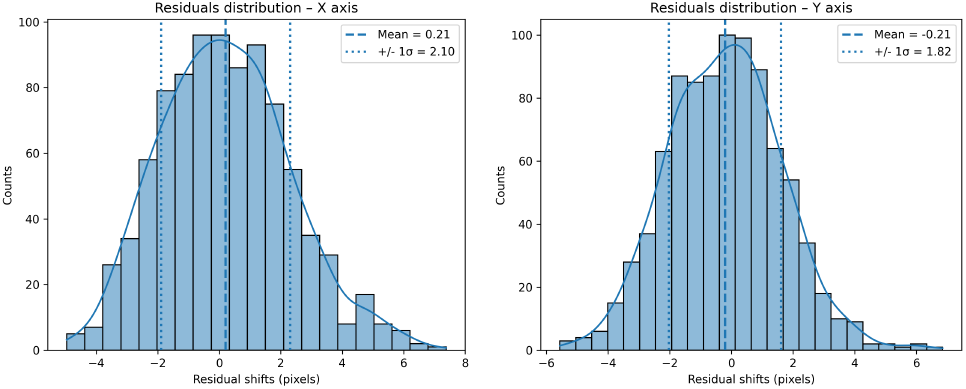
Residual shift distributions along the X and Y axes obtained from retracking experiments with imposed displacements. Histograms show the residual error, defined as the difference between the recovered shift and the known applied displacement, after correcting for the baseline tracking. Solid vertical lines indicate the empirical mean residual, while dashed lines denote the ±1*σ* interval, where the mean and standard deviation are computed directly from the residual samples using NumPy statistics (not derived from the kernel density estimate). Kernel density estimates (KDE) are overlaid to visualize the shape of the error distributions. The residuals are centered close to zero and remain small relative to the field of view, indicating stable and largely unbiased tracking performance.

### Runtime performance

The average runtime of our deep learning tracking pipeline was evaluated with and without object detection. We used a local workstation equipped with an NVIDIA RTX 3050 GPU (4 GB memory) and a baseline temporal window of 10 frames. Tests were performed on a dataset of 30 consecutive time points, and the reported runtimes represent the mean across all iterations.

Incorporating object detection marginally increased computation time. We explored the effect of reducing the image size by downsampling, which selects a subset of pixels. However, for image downsampling factors above ×2, the increase in runtime remained negligible due to the efficiency of GPU-accelerated inference. From a downsampling factor of ×4 onward, the runtime per frame dropped below one second, which is well within the temporal constraints of typical live light-sheet imaging of embryos. Memory usage remained stable, dominated by the CoTracker3 model, which required approximately 3 GB of GPU memory; the addition of the detection module introduced an overhead of about 700 MB (Tab. 1).

**Table 1.** Runtime and GPU memory usage of LiLiTTool under different down-sampling conditions. Average inference time and GPU memory consumption are reported for LiLiTTool at multiple image downsampling factors, evaluated with and without object detection. The downsampling factor *×n* denotes retention of one pixel every *n* in both the X and Y dimensions (e.g., *×*4 corresponds to one pixel retained out of four in each lateral direction), while the Z dimension was kept at full resolution. Measurements were obtained under identical hardware and acquisition conditions.

| Downsample<br>Factor | tracking-only (tracking+fusion) |  |
| --- | --- | --- |
|  | Runtime [s] | GPU Usage [GB] |
| ×1 (no scaling) | 6.13 (6.33) | 3.2 (3.9) |
| ×2 | 1.21 (1.43) | 3.3 (3.9) |
| ×4 | 0.61 (0.73) | 2.9 (3.6) |
| ×8 | 0.46 (0.63) | 2.9 (3.6) |
| ×16 | 0.43 (0.62) | 2.9 (3.6) |

Higher-resolution volumes required proportionally more GPU memory and incurred longer data transfer times, especially when performing non-local model evaluation. Consequently, downsampling proved advantageous not only for computational speed but also for mitigating data transfer overhead and maintaining compatibility with hardware constraints. For our 2304 × 2304 images, a moderate downsampling factor of ×4 therefore represented an effective balance between performance and efficiency.

### Effect of data reduction methods on tracking performance

To improve runtime efficiency and reduce data transfer overhead, several data reduction strategies were evaluated:

- **Spatial resolution reduction:** Two approaches were tested—(1) *downsampling*, which selects a subset of pixels (e.g., every 16^th^), and (2) *downscaling*, which typically involves pixel averaging or interpolation. Downsampling, implemented as array slicing, preserves large-scale spatial structures, whereas down-scaling produces smoother images and may retain greater structural consistency.
- **Bit depth reduction:** Conversion of the original `uint16` image data to `uint8` format.

To assess whether these reductions affected tracking accuracy, we conducted offline re-tracking experiments on previously acquired movies, comparing the computed stage shifts across different configurations. These shifts represent the displacement of the ROI center relative to a fixed reference position, constant across all tracking runs. Consequently, comparing the computed shifts provides a direct measure of positional consistency between methods.

Results showed minimal differences across all tested configurations: less than 3 *µ*m when comparing `uint8` and `uint16` images, and less than 6 *µ*m between downsampling and averaging. These deviations correspond to under 1% of the total FOV (~700 *µ*m), indicating negligible impact on overall tracking precision.

These findings show that all tested reduction strategies yield equivalent tracking outcomes. Given that the objective is to follow the global motion of the embryo rather than fine-scale cellular dynamics, measuring high frequency movement fluctuations is not essential. Therefore, downsampling was favored for its computational simplicity, and the `uint8` format was adopted to further optimize performance—particularly in remote inference scenarios where data transfer efficiency is critical.

### Failure cases or instabilities

Although LiLiTTool approaches demonstrated robustness, certain challenging scenarios still resulted in tracking failures or instabilities:

- **Abrupt and large embryo movements:** In one experiment, the embryo experienced a sudden and rapid rotation, causing the tracked region to move completely out of the field of view between two consecutive acquisitions. This abrupt displacement resulted in a total loss of tracking, as no valid correspondence could be established.
- **Embryo collision and occlusion:** In another case, two embryos came into close contact, leading to partial overlap within the same XY region. In contrast with the example in supplementary movie 7, one embryo appeared markedly brighter than the other, introducing a bias in the CoTracker predictions. Furthermore, the overlap generated ambiguities in the maximum-intensity projection, which compromised centring along the Z-axis resulting in the shift of the ROI from one embryo to the other.
- **Point hijacking by transient artifacts:** In a separate instance, a bright particle moved rapidly across the FOV and temporarily attracted one of the CoTracker points. While this event did not cause a complete tracking failure, it introduced a transient instability until the particle escaped CoTracker’s attention, illustrating the model’s sensitivity to unexpected bright artifacts or moving debris.

## Discussion

In this work, using zebrafish embryos imaged with a light-sheet microscope as a model, we developed and validated a real-time tracking framework for 3D biological imaging called LiLiTTool. The algorithm is designed to maintain precise positioning of features of interest—such as the tail tip—in the microscope’s FOV throughout long-term time-lapse acquisitions. By integrating deep learning–based motion estimation (CoTracker3) with an optional object detector (Faster R-CNN) and combining their outputs through a sensor fusion strategy, the system achieves accurate and stable tracking even under complex deformations and global embryo movements. The method operates within real-time acquisition constraints and is fully integrated into the Viventis LS1 microscope ecosystem, demonstrating its practicality for routine experimental use. The tool has been successfully adopted and routinely employed by members of the Timing, Oscillation, Patterns laboratory at EPFL.

### Limitations of CoTracker3

Despite its overall robustness, CoTracker3 can exhibit instability in the presence of strong, unexpected image artifacts, as described in the failure cases. For instance, a bright particle moving rapidly across the field of view may temporarily attract a tracking point, leading to local inaccuracies in the estimated motion field. One possible mitigation strategy would be to introduce a filtering mechanism that detects and removes trajectories inconsistent with the dominant motion pattern. However, defining a reliable criterion for such inconsistencies is non-trivial. Overly strict filtering risks discarding valid but atypical motion trajectories. Developing a principled and adaptive approach for discriminating between erroneous and meaningful motion deviations remains a promising direction for future work.

### Limitations of the sensor fusion strategy

The proposed sensor fusion framework effectively integrates tracking and detection-based estimates, combining their complementary strengths to enhance robustness. However, several limitations remain. In the current implementation, the tracker serves as the temporal reference against which detections are validated and fusion weights are computed. Consequently, when the tracker experiences substantial drift—such as after extended periods without reliable detections—the system may reject correct detections that appear inconsistent with the tracker’s prediction. This design choice introduces a bias toward temporal stability, potentially at the expense of responsiveness to valid corrections.

Furthermore, the weighting mechanism relies solely on spatial overlap between the predicted and detected regions, while the sigmoid function controlling the fusion parameter *α* involves two tunable hyperparameters (*k, c*) that must be adjusted manually. Overall, the method favors robustness to false positives and smooth temporal transitions rather than aggressive realignment, under the assumption that consistent detections will eventually re-synchronize the system. As a result, the reliability of the detection model becomes critical. A well-trained, task-specific detector substantially reduces the risk of consecutive detection failures, thereby maintaining effective fusion performance over long acquisitions.

### Future directions

#### Other model systems

We have demonstrated the functionality of LiLiTTool using elongating zebrafish embryos, and tested it also on other single and multi-cell systems. It will be a natural extension to deploy LiLiTTool for imaging of 3D-stem cell-based culture models that show growth and deformation similar to that of embryos, such as gastruloids, somitoids or trunk-like structures (29), or organoids with growing villi, branches or other protrusions (30).

#### Online Applications

Although the current implementation performs reliably, several avenues for improvement remain. Implementing asynchronous tracking and stage repositioning could decouple computation time from acquisition intervals, thereby improving resilience to unexpected processing delays. Expanding the detection model to include multiple anatomical landmarks or targets would broaden the system’s applicability and enable multi-region tracking. The incorporation of multiple, shape-aware ROIs could also enhance robustness by allowing the system to track both position and orientation, ensuring that the maximal portion of the target structure remains within the field of view.

A further direction involves extending CoTracker3 to estimate motion along the Z-axis, enabling selective tracking of individual planes within the volume rather than the entire embryo. This would improve accuracy while reducing computational and storage costs. Additionally, adapting the pipeline for real-time cell-level tracking could open new opportunities in live-cell imaging applications, such as optogenetic stimulation (31) or microsurgical manipulation guided by dynamic feedback (32).

Finally, establishing a benchmark dataset and standardized evaluation metrics for zebrafish embryo tracking would facilitate objective performance evaluation and promote reproducibility and methodological innovation within the field.

#### Offline Applications

Beyond real-time operation, the proposed framework also offers potential for offline analysis. For instance, it can be adapted for cell tracking on pre-acquired time-lapse datasets, enabling detailed reconstruction of cell trajectories and behaviors. Moreover, CoTracker3 supports back-tracking, allowing the identification of earlier positions of cells or cell groups that become relevant at specific developmental stages. Such offline analyses could help elucidate lineage relationships or spatial organization processes.

## Materials and Methods

### Zebrafish husbandry and embryos

Zebrafish were raised under standard conditions (33). Adult zebrafish (*Danio rerio*) were maintained under standard laboratory conditions in a recirculating system at 26°C, with a 14h/10h light/dark cycle. Water quality was maintained at a pH of 7.5 and conductivity of 600*µ*S. Embryos were obtained from natural spawning and maintained in system water at 28.5°C. We used an in-cross of heterozygous genotypes, H2bmCherry and selected a mix of heterozygous and homozygous embryos for the experiments to enable varying of signal intensity. Embryos were prepared for microscopy following the methods described in (17). Heat-shock experiments delivered a pulse of elevated temperature (40°C for 30 minutes) using a peristaltic pump (Grothen model G728-2) to circulate water through a waterbath (Isotemp™ model FSGPD05) and into the light-sheet microscope sample chamber. All procedures involving zebrafish were performed in accordance with the guidelines approved by the federal food safety and veterinary office of the canton of Vaud, Switzerland (authorization no. VD-H23).

### Microscope choice

All experiments were conducted using the LS1 Live light-sheet microscope (Viventis GmbH, Switzerland, https://www.leica-microsystems.com/products/p/viventis-ls1-live/). The Viventis LS1 Live was designed for high-resolution, long-term imaging of large three-dimensional biological specimens under controlled physiological conditions. Its dual-illumination and single-detection architecture ensures high image quality, while the open-top design facilitates straightforward specimen mounting and allows for medium exchange during time-lapse acquisitions. Furthermore, its multi-position imaging capability and customizable acquisition parameters at each position provide experimental flexibility. Open access to the Viventis PyMCS control software was essential for closed-loop microscopy.

### Microscope performance assessment

The LS1 system employs a high-precision Cartesian linear stage capable of movement along the X, Y, and Z axes. Accurate stage positioning is essential for reliable imaging and automated tracking, particularly during extended time-lapse acquisitions. To verify the positioning precision and repeatability of the microscope, we conducted a systematic performance assessment under controlled motion conditions.

Fluorescent beads embedded in agarose gel were used as stable, immobile reference markers to quantify stage motion. This choice ensured a stationary and easily segmentable reference object, enabling robust measurement of true displacement. A scripted stage motion sequence was executed, and corresponding images were recorded to assess both accuracy and repeatability.

The analysis pipeline involved converting bead images into 3D point clouds, with each bead represented as a single point. The resulting point clouds were aligned using the Iterative Closest Point (ICP) algorithm (34) implemented in the `Open3D` library (35). The evaluation demonstrated sub-pixel accuracy in the XY plane and sub-step precision along the Z-axis, confirming that the microscope meets the required specifications for closed-loop tracking.

### LiLiTTool performance assessment without ground truth

Quantitative evaluation of tracking accuracy is inherently challenging for long-term, live imaging of zebrafish embryos, not only because explicit ground truth annotations of tail motion are unavailable, but also because the task of maintaining a specific anatomical feature within the field of view does not admit a unique ground truth trajectory. Indeed, multiple tracking solutions can be equally valid for this objective, and manual annotations would likely vary across observers. To address this ambiguity, we adopted a self-consistency–based evaluation strategy that enables quantitative assessment of tracking performance under controlled and known perturbations.

Specifically, we reprocessed previously acquired and successfully tracked time-lapse datasets by introducing a known artificial displacement. For each sequence of images, the original 3D image volumes were embedded into a larger volume filled with empty pixels, enabling the application of a predefined spatial shift without introducing boundary artifacts. The modified image sequences were then re-tracked using the same *tracking-only* pipeline.

Two tracking runs were performed for each dataset: (i) a reference run on the unmodified image sequence, and (ii) a perturbed run in which a known shift was applied to the images prior to tracking both in X and Y directions. The residual error was computed as the difference between the shift estimated in the perturbed run, corrected for the known artificial displacement, and the shift estimated in the reference run. Formally, the residual shift Δ_res_ is defined as:

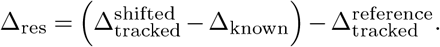

This residual reflects the tracking error introduced by the pipeline, independent of the imposed displacement. Under ideal tracking performance, the residual shift would be zero. Deviations from zero therefore provide a direct measure of the method’s sensitivity to known perturbations and its robustness to image translations.

The residual shifts were quantified simultaneously along the X and Y image axes. Across all evaluated configurations, the mean residual shift was +0.21 pixels in X and −0.21 pixels in Y, indicating negligible systematic bias in the tracking estimates. The corresponding standard deviations of the residuals were 2.1 pixels in X and 1.82 pixels in Y, reflecting the variability of the tracking error under imposed perturbations (Fig. 4). Residuals along the Z axis are not reported. Unlike lateral (X–Y) Cotracker3 tracking, axial position is estimated using a center-of-mass–based measurement on a binary representation of the structure of interest. As long as the tracked feature remains fully contained within the field of view along Z, this estimate is invariant to small axial translations of the image volume. Consequently, re-tracking experiments with imposed Z shifts yield negligible or ill-defined residuals that do not reflect tracking accuracy but rather the intrinsic property of the estimator. For this reason, only X and Y residuals, which directly quantify lateral tracking precision and stage correction performance, are reported.

The residuals remains small relative to the microscope field of view (2304 pixels) and the characteristic scale of tail motion. Overall, these results demonstrate that the tracking pipeline accurately recovers imposed displacements and maintains stable performance in the absence of ground truth annotations.

While this approach does not replace true ground truth validation, it provides a practical and reproducible means to assess tracking accuracy in scenarios where manual annotation is infeasible and directly probes the consistency and stability of the tracking algorithm under controlled conditions.

### Tracking algorithms considered

Three complementary tracking strategies were evaluated:

1. **Cross-correlation-based tracking:** This classical method computes voxel-wise similarity between consecutive frames in Fourier space, achieving sub-pixel precision by upsampling correlation peaks. Although robust under simple conditions, it performed poorly under significant embryo deformations or rotations and was computationally expensive. Consequently, it was not retained for further use.
2. **Deep learning based tracking (CoTracker3):** CoTracker3 is a transformer-based model designed for dense 2D point tracking in video sequences. We adapted it to microscopy data by applying it to maximum intensity projections and tracking a dense grid of query points in the XY plane. This model captures non-linear motion and remains robust to local intensity variations.
3. **Sensor fusion with Faster R-CNN detection:** A detection model fine-tuned on zebrafish embryo tail tips provides independent observations, which are integrated with the tracking estimates using a Kalman-like fusion framework. This combination corrects drift and dynamically adapts the ROI dimensions during acquisition.

**Deep learning based tracking with CoTracker3**. Because CoTracker3 operates exclusively in 2D, we use it to estimate motion in the XY plane from maximum intensity projections as follows:

1. **Preprocessing:** Images are first downsampled (typically by a factor of four) to improve processing speed, and maximum intensity projections are computed along Z to generate 2D representations. A sliding temporal window is constructed so that, for each time point *t*, a short video sequence

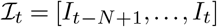

is formed, with the last frame corresponding to the current image. Normalization is performed using the 1st and 99th intensity percentiles as scaling bounds:

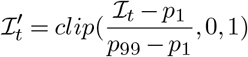

which ensures consistent brightness and contrast while reducing the impact of outlier intensities.
Original microscopy images are acquired in 16-bit format (`uint16`), but are converted to 8-bit (`uint8`) for efficiency, particularly in remote inference settings. Despite the loss of dynamic range, this conversion yielded no measurable degradation in tracking accuracy. Query points are generated by binarizing the ROI mask and sampling a uniform grid of points within it.
2. **Model inference:** The normalized video and query points are passed to CoTracker3, which predicts the positions of the tracked points over time, denoted [*T*_*t*−*N*+1_, …, *T*_*t*_].
3. **ROI update:** To infer global XY motion, displacement vectors between corresponding points at *t* − 1 and *t* are computed as *V* = *T*_*t*_ − *T*_*t*−1_. A linear regression model is then fitted to these displacements to estimate translation and linear deformation:

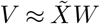

where 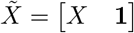 contains the tracked coordinates at *t* − 1, and *W* encapsulates the affine transformation and translation components. The optimal parameters are obtained via least squares using `NumPy`’s `linalg.lstsq` solver. The ROI center is subsequently updated using the predicted affine transformation.

### Estimate position in Z

Z tracking aims to maintain the full structure within the imaging volume. The Z displacement is computed separately using a classical shape-dependent method, motivated by the observation that embryo deformation and motion are more complex in XY than in Z. Overlaps between body parts can occur in XY projections but are rare along the Z-axis, where structures remain simpler and motion amplitude is lower. The image is cropped using the updated XY ROI and binarized via Otsu’s (20). method. The Z position is then estimated as the intensity-weighted center of mass:

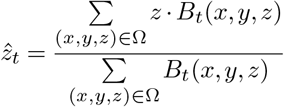

where *B*_*t*_ is the binary mask and Ω the ROI voxel set.

### Object detection model

Although CoTracker3 provides accurate motion estimation, two key limitations were identified: (i) gradual drift accumulation due to purely predictive updates, and (ii) fixed ROI size and shape, which can become misaligned during embryo deformation. To address these issues, a detection model provides independent frame-by-frame estimates of the target region (typically the tail tip). The detection network is based on the `torchvision` implementation of Faster R-CNN v2 (25, 26) with a ResNet-50 backbone and Feature Pyramid Network (FPN). The FPN captures multi-scale image features, while the Region Proposal Network (RPN) generates candidate bounding boxes subsequently refined and classified by the Fast R-CNN head. This detection model was fine-tuned for 100 epochs on a task-specific dataset of 400 2D images, each annotated with a bounding box around the zebrafish embryo tail tip. Data augmentation included random rotations, flips, and contrast variations. As a result, the model is specialized for this anatomical feature and would require re-training for other targets.

### Sensor fusion of CoTracker3 and detection model

The use of an object detection model aligns naturally with a sensor fusion framework, in which CoTracker3 serves as a predictive model and the Faster R-CNN detection acts as an observation model.

While classical Kalman filtering (21) assumes linear Gaussian noise, the deep neural network–based detector may produce non-Gaussian errors or false positives. Therefore, a heuristic fusion strategy was adopted, using geometric consistency as a proxy for detection reliability. Fusion proceeds in two steps:

1. **Detection validation:** A feature detection is accepted based on the following conditions:

- *Intersection ratio:* The overlap between tracker and detector ROIs, normalized by the smaller ROI’s area.
- *Size ratio:* The ratio of largest ROI relative to smallest ROI. This ensures the detected region is not unrealistically large.
- *Detection score:* The detector’s softmax score, used in combination with the above metrics.
2. **Fusion weighting:** The fusion weight *α* is derived from the intersection ratio (*r*) via a sigmoid function:

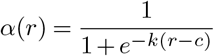

where *c* defines the inflection point and *k* the slope. When tracker and detector are aligned, the tracker dominates; when misaligned, the detector correction prevails; and if validation fails, the detection is ignored (*α* = 0).

The detection-derived bounding box also dynamically updates ROI dimensions, compensating for embryo rotation and deformation. Fusion is performed in XY, while Z tracking remains independent.

### Tracking modes

LiLiTTool provides several tracking modes to accommodate different experimental needs and prioritization strategies. In SingleROI mode, a single region of interest (ROI) is defined and tracked, and stage movements are computed exclusively to maintain this ROI within the field of view (FOV). This mode is intended for experiments focusing on a single anatomical feature.

For experiments involving multiple regions, LiLiTTool offers three MultiROI modes. In MultiROI – max ROIs (non-weighted) mode, stage updates are computed to maximize the number of ROIs retained within the FOV, without prioritizing any specific ROI. All ROIs contribute equally to the stage correction estimate.

In MultiROI – max ROIs (weighted) mode, the objective remains to retain as many ROIs as possible within the FOV, but individual ROIs are assigned weights according to their userdefined order. This allows preferential retention of selected ROIs when trade-offs are required.

Finally, MultiROI – priority mode is designed to maintain all ROIs within the FOV while strongly prioritizing the first ROI in the user-defined list. In this configuration, stage updates are primarily driven by the highest-priority ROI, while secondary ROIs are incorporated with lower influence. This mode is particularly suited for experiments in which one anatomical structure is critical, but additional regions should be preserved whenever feasible.

### LS1 system integration

Direct hardware communication is unavailable during acquisition on the LS1 system. Instead, synchronization is achieved through the microscope’s custom API (*PyMCS*), which emits a signal after each acquisition. Upon receiving the signal, the tracking script pauses the time-lapse, processes the newly acquired image, updates the ROI, computes new stage coordinates, and sends them back to the microscope before resuming acquisition.

This synchronous design guarantees adaptive repositioning but imposes strict timing constraints: the tracking computation must complete before the next acquisition begins. To ensure reliability, the algorithm moves the stage to the position estimated from the previous frame (*t* − 1), avoiding speculative motion prediction. An asynchronous tracking mechanism could further enhance robustness by decoupling computation from acquisition timing, and is compatible with existing Viventis workflows.

### Remote inference GPU execution

Since the local acquisition machines lack compatible GPUs, model inference is offloaded to a remote GPU server via the `imaging-server-kit` framework. This client–server setup allows the client to preprocess images, transmit data, and receive model outputs from the remote GPU.

CoTracker3 and Faster R-CNN together require approximately 3.5 GB of GPU memory. CoTracker3 processes a temporal window of 10 downsampled frames, while Faster R-CNN operates on the most recent 2D projection.

Robust fallback mechanisms handle network or server interruptions to prevent acquisition deadlocks: if inference fails, the system skips the iteration and resumes normal operation until the connection is restored.

### Code architecture and design

The software architecture follows a modular and extensible design, enabling both real-time interaction with microscope hardware and offline analysis of pre-recorded datasets. At its core, the **TrackingRunner** module coordinates all major components, including the **PositionTracker, ROITracker, MicroscopeInterface** (Fig. 5).

**Fig. 5.**
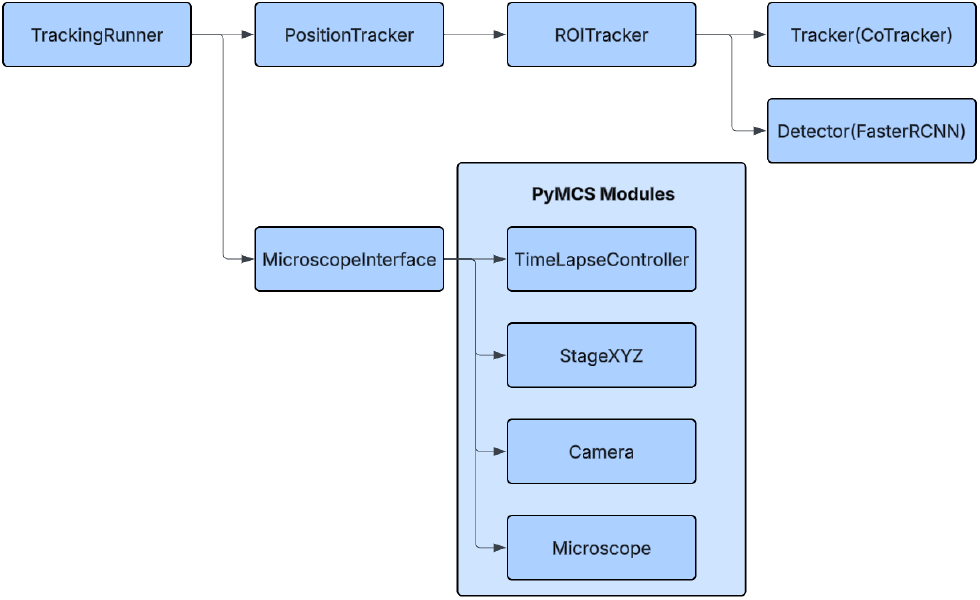
Modular software architecture of LiLiTTool and interactions between its core components. The *TrackingRunner* orchestrates the tracking pipeline and coordinates communication between the *PositionTracker, ROITracker*, and *MicroscopeInterface*. The *PositionTracker* converts image-based displacements into physical stage movements, while the *ROITracker* updates regions of interest by combining deep learning–based motion estimation (CoTracker3) with optional object detection (Faster R-CNN). The *MicroscopeInterface* provides an abstraction layer for hardware control and image access; a dedicated implementation for the Viventis LS1 microscope is shown (*PyMCS modules*). This modular design enables multi-embryo tracking, supports both real-time and offline workflows, and facilitates adaptation to alternative microscope systems and acquisition pipelines.

The **PositionTracker** manages embryo-level tracking by computing displacements and translating pixel shifts into stage movements. The **ROITracker** combines outputs from the deep learning tracker (CoTracker3) and the object detection model (Faster R-CNN) to update the region of interest. The **MicroscopeInterface** provides an abstraction layer to communicate with the microscope control software (default Viventis PyMCS), while remaining adaptable to other hard-ware APIs. It also emcompass a generalized image access method, allowing compatibility with different image formats and acquisition schemes.

This modular organization ensures clear separation of concerns, facilitates maintenance, and supports multi-embryo tracking through independent **PositionTracker** instances. Moreover, the architecture’s interchangeable module design enables straightforward adaptation to new microscope systems or data pipelines, reinforcing its scalability and long-term usability.

### User interface and visualization

Several user interfaces were developed to support usability, monitoring, and debugging, primarily based on the `bokeh` framework (36) and integrated into the Viventis software environment.

- **ROI and detection configuration:** Enables manual selection of initial ROIs and testing of detection models performance prior to experiment start.
- **Viventis LS1 setup interface:** Built upon the *PyMCS* GUI, this interface allows users to configure pixel and step sizes, output directories, remote inference options, and operation mode (live or simulated).
- **Monitoring Tool:** Provides real-time visualization of tracking results, including ROI overlays, query points, stage displacements, and cumulative trajectories.
- **Annotation tool:** Allows manual labeling of regions for custom detector fine-tuning.
- **Offline tracking dashboard:** Enables manual point selection and bidirectional tracking across selected time windows for retrospective analysis.
- **Napari plug-in:** Offers offline tracking functionality equivalent to the standalone interfaces within the Napari ecosystem.

## ACKNOWLEDGEMENTS

We thank the EPFL Center for Imaging and the Timing, Oscillation, Patterns Lab (EPFL) for support, and in particular contributions from Mallory Wittwer, Feyza Arslan and Agnese Fronte. We would like to thank the EPFL fish facility staff for animal care. We thank Jasper Phelps and Pavan Ramdya (EPFL) for the tardigrade dataset and Coralie Busso and Pierre Gönczy (EPFL) for the *Caenorhabditis elegans* dataset.

## Data and software availability

The source code produced and used in this work can be found in GitHub https://github.com/EPFL-TOP/lightsheet-live-tracking-tool. Data can be found in the EPFL public Data Repository.

## Contribution

Conceptualization: CH, EA, FA; Methodology: FP, CH, EA, FA; Software Developments: FP, CH, FA; Supervision: CH, EA, FA; Funding acquisition: AO; Resources: FP, CH, EA, FA; Investigation: FP, CH, EA, FA; Project administration: EA, AO; Writing – original draft: CH, FP; Writing – review & editing: CH, FP, AO, J-YT, EA, FA, EV

---

**Supplementary Note 1: Supplementary figure 1**

**Query point generation process**(a) A Gaussian filter is applied to the input image to reduce local intensity variation within the tail region, helping to stabilize values before segmentation. (b) Otsu’s thresholding method is used to generate a binary mask of the tail. (c) A uniform grid of points is generated, and the mask is used to filter points outside the detected tail region. (d) The final query points are constrained by both the binary mask and the manually defined ROI.

**Supplementary Note 2: Supplementary movie 1**

**Zebrafish embryo imaged without tracking** This movie shows a real-time 30-hour acquisition without tracking enabled. The zebrafish tail tip quickly goes out of the field of view.

**Supplementary Note 3: Supplementary movie 2**

**Zebrafish embryo imaged with *tracking-only*** This movie shows the same embryo as *Supplementary movie 1* but with *tracking-only* enabled. The zebrafish tail tip also eventually goes out of the field of view due to the tracker following internal movements.

**Supplementary Note 4: Supplementary movie 3**

**zebrafish embryo imaged with *tracking+fusion*** This movie shows the same embryo as *Supplementary movie 1* but with *tracking+fusion* enabled. The zebrafish tail tip is perfectly tracked.

**Supplementary Note 5: Supplementary movie 4**

**Zebrafish embryo retracking comparison:** This movie shows an in silico retracking example that compares expert human tracking with the various tracking options offered by LiLiTTool.

**Supplementary Note 6: Supplementary movie 5**

**Zebrafish embryo imaged with auto-fluoresence:** This movie shows zebrafish embryos imaged in low fluorescence conditions, relying only on the auto-fluorescence of the fish.

**Supplementary Note 7: Supplementary movie 6**

**Zebrafish embryo imaged during heat-shock experiment:** During such conditions, the embryo undergoes a rapid temperature increase, leading to contractions and sudden large displacements.

**Supplementary Note 8: Supplementary movie 7**

**Zebrafish embryo imaged with another in the FOV:** LiLiTTool accurately follows the selected ROI (*tracking-only no blurring*).

**Supplementary Note 9: Supplementary movie 8**

***Tardigrades* time-laps:** LiLiTTool accurately tracks the position of the animal’s head across time, maintaining stability despite strong deformations and non-rigid movements of the organism.

**Supplementary Note 10: Supplementary movie 9**

**Early *Caenorhabditis elegans*embryos imaged during mitosis:** the method accurately followed the fluorescently-labelled spindle poles during cell division, capturing their fast and non-linear trajectories within a crowded cellular environment.

**Supplementary Note 11: Supplementary movie 10**

**Human hepatocarcinoma-derived (HepG2) cells from the Cell Tracking Challenge benchmark datasets:** The system successfully tracked 17 individual HepG2 cells independently throughout the sequence.

