## Supplemental Figure 1 for "Real-Time Multi-Position and Multi-ROI Tracking with LiLiTTool for Smart Light-Sheet Microscopy in Growing Samples"

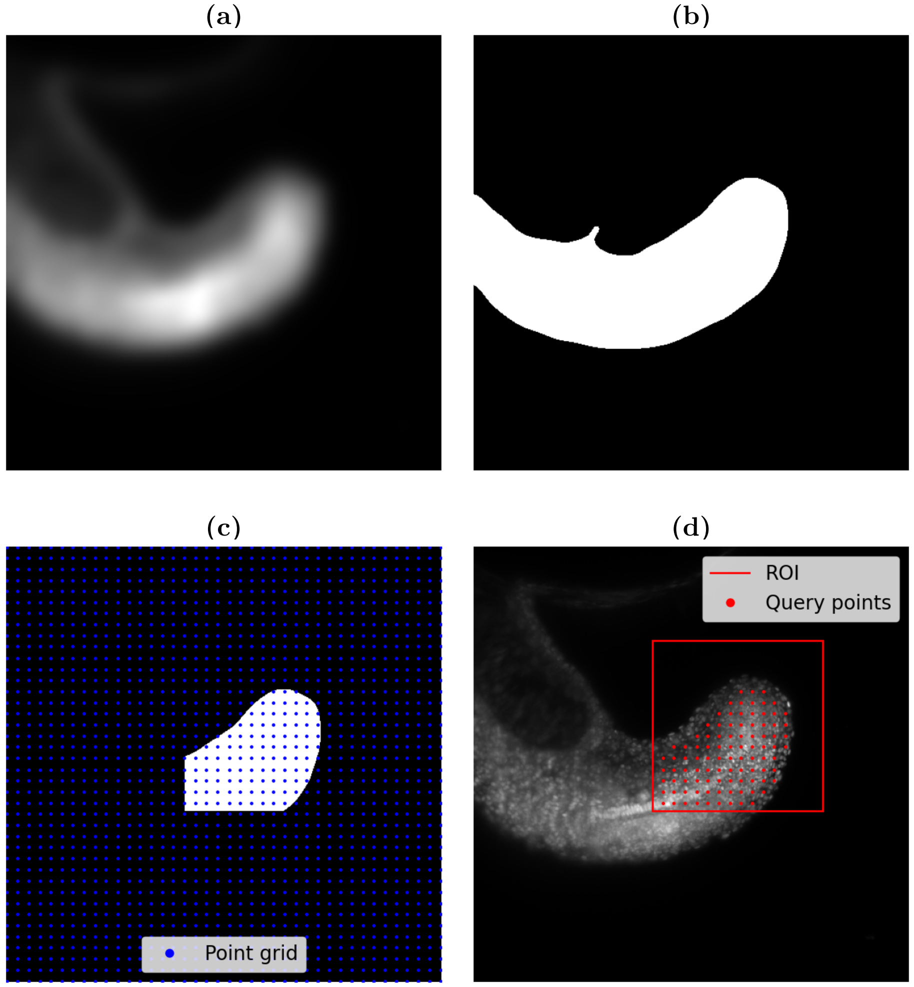


Supplemental Figure 1: **Query point generation process.**
**(a)** A Gaussian filter is applied to the input image to reduce local intensity variation within the tail region, helping to stabilize values before segmentation. **(b)** Otsu's thresholding method is used to generate a binary mask of the tail. **(c)** A uniform grid of points is generated, and the mask is used to filter points outside the detected tail region. **(d)** The final query points are constrained by both the binary mask and the manually defined ROI.
